# A key regulatory protein for flagellum length control in stable flagella

**DOI:** 10.1101/872952

**Authors:** Madison Atkins, Jiří Týč, Shahaan Shafiq, Manu Ahmed, Eloïse Bertiaux, Artur Leonel De Castro Neto, Jack Sunter, Philippe Bastin, Samuel Dean, Sue Vaughan

**Affiliations:** Biological & Medical Sciences, Oxford Brookes University, Gipsy Lane, Oxford, OX3 0BP, UK; Trypanosome Cell Biology Unit and Inserm U1201, Institut Pasteur, 25, rue du Docteur Roux, 75015, Paris; Warwick Medical School, University of Warwick, Coventry, CV4 7AL, UK

**Keywords:** Cep164, ciliogenesis, flagella, cilia, Trypanosoma, basal body, transition fibres

## Abstract

Cilia and flagella are highly conserved microtubule-based organelles that have important roles in cell motility and sensing [1]. They can be highly dynamic and short lived such as primary cilia or *Chlamydomonas* [2] or very stable and long lived such as those in spermatozoa [3] photoreceptors [4] or the flagella of many protist cells [3,4]. Although there is a wide variation in length between cell types, there is generally a defined length for a given cell type [1]. Many unicellular flagellated and ciliated organisms have an additional challenge as they must maintain flagella/cilia at a defined length whilst also growing new flagella/cilia in the same cell. It is not currently understood how this is achieved. A grow-and-lock model was proposed for the maintenance of stable flagella where a molecular lock is applied to prevent flagellum length change after assembly [5]. The molecular mechanisms of how this lock operates are unknown, but could be important in cells where an existing flagellum must be maintained whilst a new flagellum assembles. Here we show that Cep164C contributes to the locking mechanism at the base of the flagellum in *Trypanosoma brucei*. It is only localised on the transition fibres of basal bodies of fully assembled flagella and missing from assembling flagella. In fact, basal bodies only acquire Cep164C in the third cell cycle after they assemble in trypanosomes. Depletion leads to dysregulation of flagellum growth with both longer and shorter flagella; consistent with defects in a flagellum locking mechanism. By controlling delivery of components into the old assembled flagellum, maintenance of stable flagella can occur but limits further growth. This offers an important explanation for how many eukaryotic unicellular cells maintain their existing flagella whilst growing new ones before these cells divide. This work also reveals additional regulatory roles for Cep164 in eukaryotic organisms.

## Results and discussion

### Cep164C acquired in the third cell cycle after basal body formation

The boundary between the cytosol and the flagellum where the transition fibres and transition zone are located is proposed to have functions in allowing selective targeting of molecules into the flagellum [6]. Trypanosomes are pathogenic protists whose single flagellum remains assembled during the cell cycle with a new flagellum assembled alongside during every cell division [7]. This provides an excellent model to study differential regulation of flagellum growth in a single cell where the assembled old flagellum is maintained whilst a new flagellum assembles. Recent evidence suggests that the assembled flagella are prevented from further elongation via a lock mechanism [5]. We searched the TrypTag protein localisation project for proteins that localised to the old flagellum only [8] and Cep164C was identified for further studies. The Centrosome Protein (CEP) 164 is located at the distal appendages of centrioles in mammalian cells and is important for the docking of centrioles to the plasma membrane for assembly of cilia [9–11]. Given its localisation, a plausible functional role could include regulating entry of components into a flagellum. A Cep164C:mNeonGreen fusion protein was expressed from the endogenous locus and co-localised with an antibody to transition fibre protein retinitis pigmentosa-2 [12]. Localisation of Cep164:mNG was observed at the proximal end of the old flagellum only and not the new flagellum in both detergent-extracted cytoskeletons and whole cells. Localisation was identical when mNG was added to the N- or C-terminus (Fig 1A-D). In dividing cells with two flagella, there is one old flagellum which was assembled in a previous cell cycle and one growing new flagellum. Cep164C was only located on the mature basal body of the assembled old flagellum (Fig 1C-D; arrow) and not the mature basal body of the new flagellum (Fig 1B-D; arrowhead), even though both mature basal bodies were docked to the plasma membrane. Cep164C remained on the old mature basal body of the assembled flagellum so at cytokinesis the daughter cell with the old flagellum contains Cep164C, but the daughter cell with the new flagellum does not (Fig 1D). Accordingly, there should be at least 50% of G1 cells positive for Cep164C and 50% negative for Cep164C (Fig 1A). However, multiple experiments revealed no more than 40%, of G1 cells with Cep164C, suggesting removal of Cep64C from the old flagellum daughter cell after cytokinesis and re-appearance at the start of the next cell cycle or that daughter cells might re-enter the cell cycle at different rates (Fig 1E; N= 800 cells).

**Figure 1:**
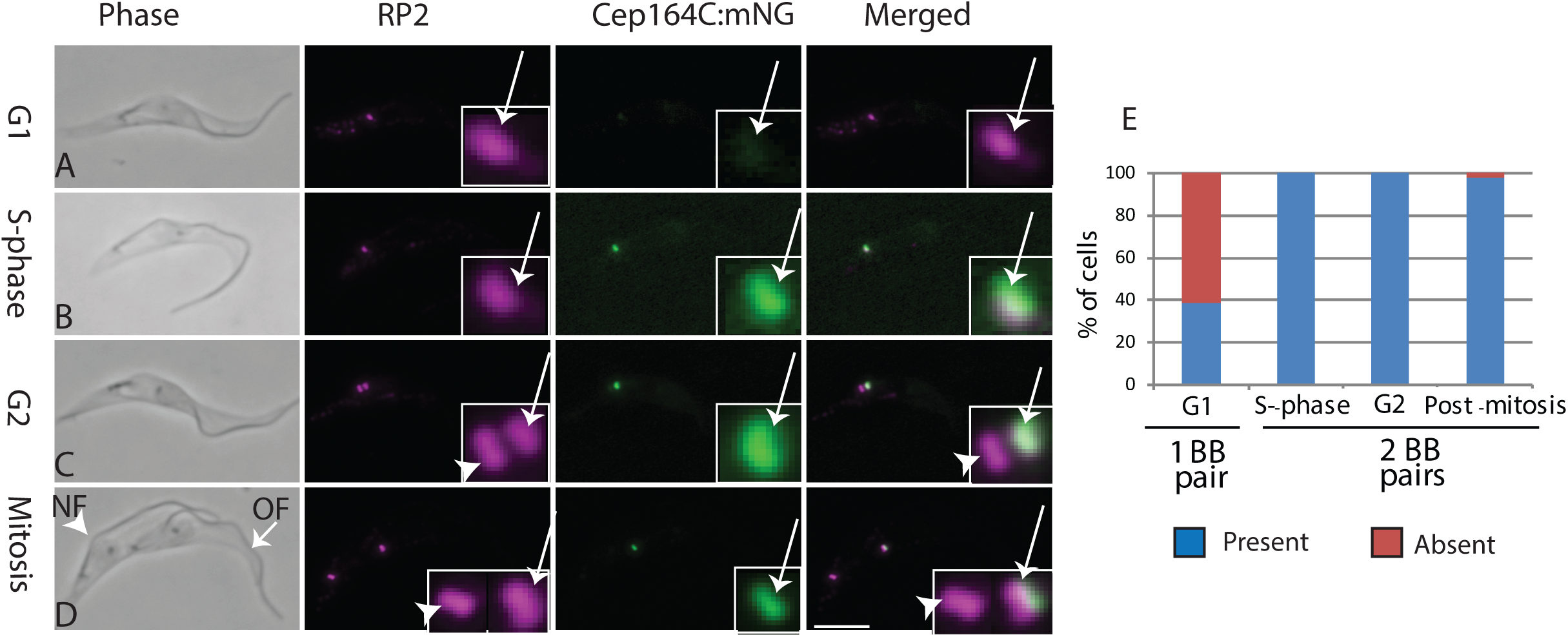
Cell cycle-dependent localisation of Cep164C; A-D: Endogenously tagged Cep164C:mNG (green) co-localised to transition fibre protein RP2 (magenta) using YL1/2 antibody in detergent-extracted cytoskeletons. Cep164C::mNG is only present on old flagellum mature basal body (arrows) and not new flagellum mature basal body (arrowheads) in dividing cells with 2 flagella. OF (old flagellum), NF (new flagellum); Scale bar = 5μm; E: cell cycle analysis of presence (blue) and absence (red) of Cep164C through the cell cycle N=200 cells per cell cycle stage.

Basal bodies and centrioles have a defined maturation lineage across multiple cell cycles and these results reveal a novel localisation pattern that distinguishes the two mature basal bodies in a single cell. After initial formation in one cell cycle, a pro-basal body/pro-centriole must proceed through a second cell cycle to mature and assemble a flagellum/cilium. In trypanosomes, it is only in the third cell cycle that Cep164C is acquired, after completion of flagellum assembly (Fig 4A,B).

**Figure 2:**
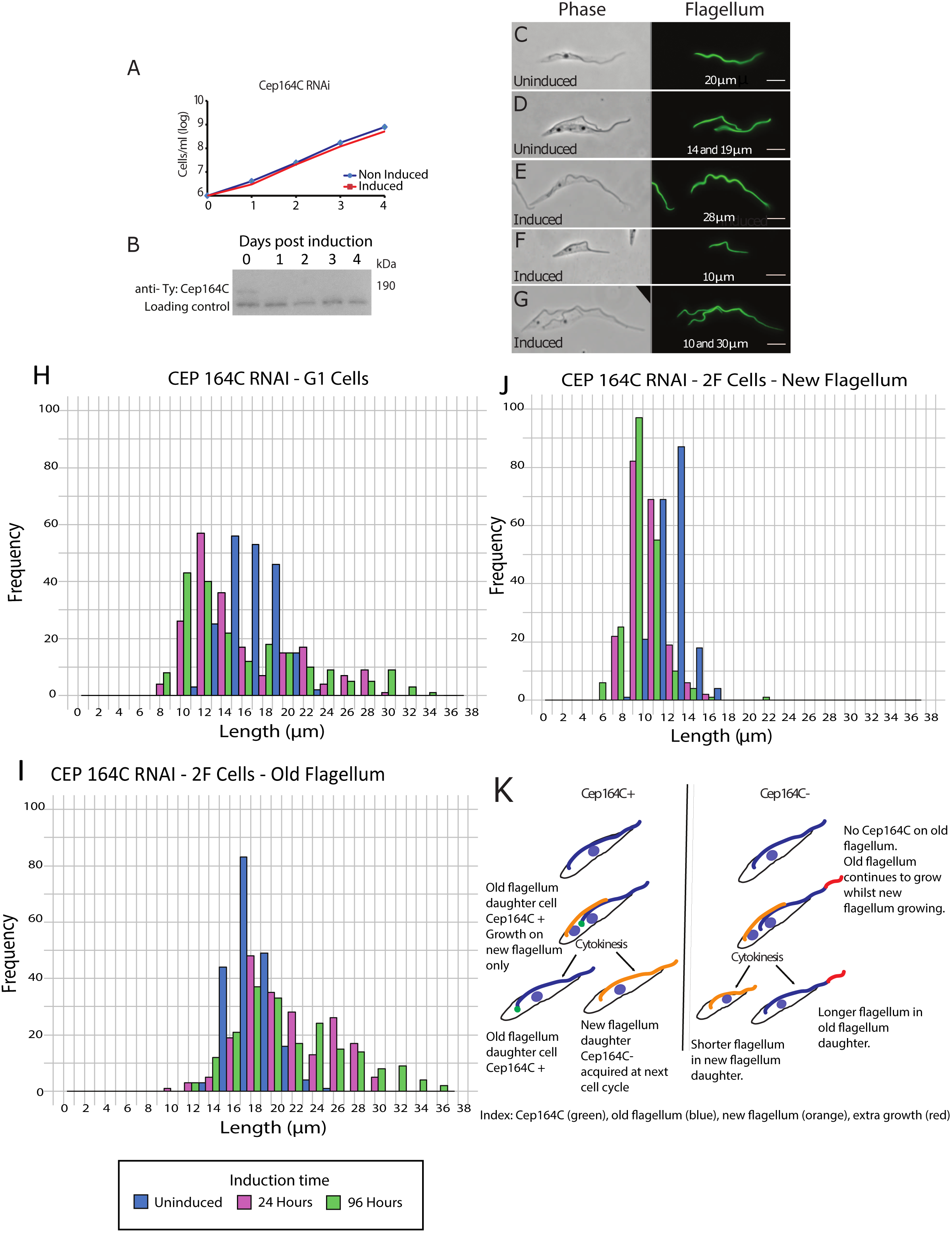
RNAi against Cep164C; A: growth curve of uninduced and induced *CEP164C*^*RNAi*^ cells; B: Western blot using anti-Ty1 antibody (BB2) to show knockdown of Cep164C over 0-4 days. This epitope flanks the mNeonGreen Cep164C and is detected using Anti-Ty (BB2) antibody; C-G: Detergent-extracted cells labelled with the L8C4 antibody (flagellum). Uninduced cells (C, D) induced (E, F,G) illustrating wide variation in flagellum lengths post induction of *CEP164C*^*RNAi*^; H-I: Measurements of flagella in uninduced and *CEP164C*^*RNAi*^ 24h and 96h post-inudction of RNAi; Single flagellum G1 cells (H), post-mitotic dividing 2F cells old flagellum (I) – new flagellum (J) - Uninduced (blue), 24hrs (red), 96hrs (green); K: Model illustrating role of Cep164C in the regulation of flagellum growth between old and new flagella.

**Figure 3:**
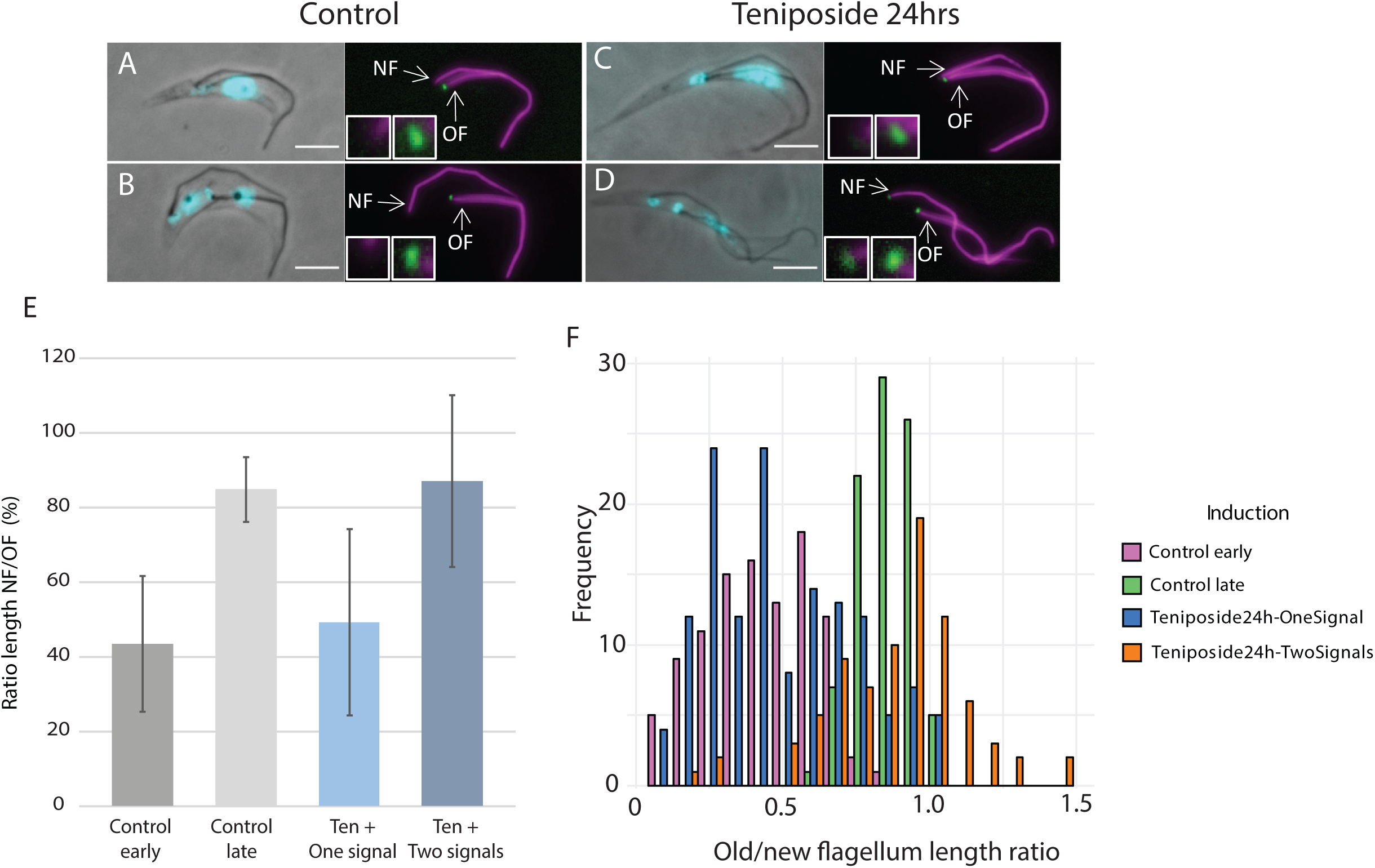
Cep164C accumulates on locked new flagella following teniposide treatment. A,B: Control dividing cells with endogenously tagged Cep164C:mNG on old flagellum mature basal body only (control; green), flagella labelled with anti-axoneme antibody Mab25 (magenta); C-D: teniposide treated cells 24hrs, C: Cep164C::mNG is only found on the old flagellum (one signal), D: Cep164C on old and new flagella (two signals). All detergent-extracted cytoskeletons; E: average ratio of old versus new flagellum lengths in control cells (early = G2 cells; late = post-mitotic cells - grey). Teniposide treated cells (Ten +) blue. The ratio of old vs new flagellum length in cells with one Cep164C signal (Ten + one signal) is less than in cells with two Cep164C signals (Ten + two signals); F: frequency histogram of old/new flagellum length ratios shown in E to illustrate spread of the data.

**Figure 4:**
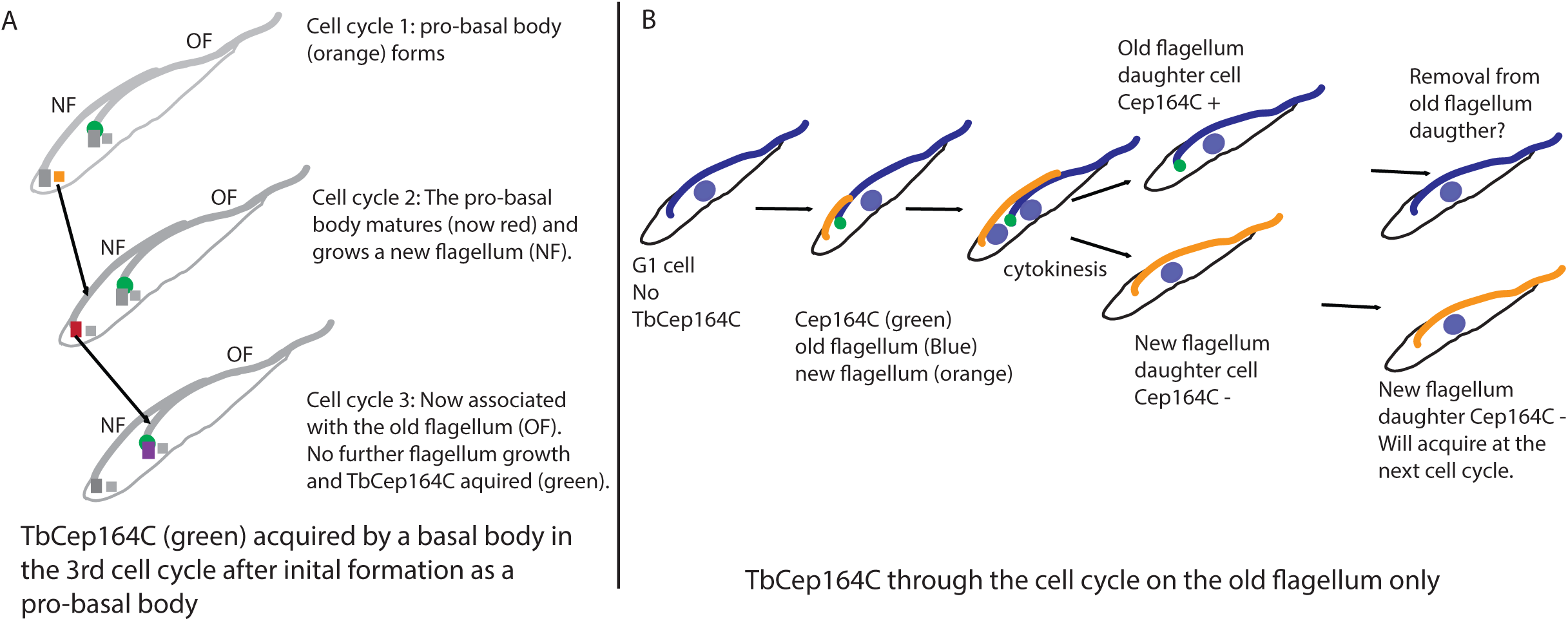
A: model illustrating acquisition of Cep164C is in the 3^rd^ cell cycle after initial formation as a pro-basal body (cell cycle 1 - orange), followed by maturation and growth of a new flagellum (cell cycle 2 - red) and finally acquisition of Cep164C in cell cycle 3 (purple) when is it associated with the old flagellum basal body and no further growth of the old flagellum occurs; B: model illustrating cell cycle dependent acquisition of Cep164C (green) onto the old flagellum (blue) and not the new flagellum (orange).

### Inducible knockdown of Cep164C leads to dysregulation of flagellum growth

We hypothesised that if Cep164C does play a role in locking further growth of the assembled old flagellum then depletion would result in dysregulation of flagellum growth on both the old and new flagella. Knockdown of Cep164C was carried out following stable transformation of trypanosomes with a plasmid expressing double-stranded RNA under tetracycline-inducible promotor control [13]. Inducible RNAi knockdown of Cep164C did not inhibit population doubling, there was no visible defect in motility and knockdown of protein expression was confirmed by western blotting (Fig 2A, B). However, light microscopy demonstrated abnormalities in the length of flagella. In cells with a single flagellum some had excessively long flagella (Fig 2E) or flagella that were too short (Fig 2F) compared to controls (Fig 2C). Measurements of flagellum length in cells confirmed these observations. Measurements of cells with a single flagellum (G1 cells) revealed significantly wider variation post-induction of RNAi (24hrs – magenta; 96hrs green) compared to uninduced (blue) (F-Test and Wilcoxon test Table S1). In post-mitotic dividing cells, it is known that the new flagellum is slightly shorter than the old flagellum at cytokinesis and final growth is completed following cytokinesis [14]. However, in the RNAi population there was a greater disparity between new and old flagellum lengths (Fig 2G) compared to controls in post-mitotic cells (Fig 2D). In these dividing cells the very long flagella were always the old flagellum whereas the short flagella were always the new flagellum. Measurements of old (Fig 2I) and new flagella (Fig 2J) in post-mitotic dividing cells showed the old flagellum could reach up to 37μm in length compared to the old flagellum in uninduced cells, which was never longer than 26 μm. The variation in flagellum lengths between uninduced and 24 or 96hrs post-induction were statistically significant (Table S1). This confirms that the wide disparity in flagellum lengths seen in G1 cells (Fig 2E; 28 μm, F; 10 μm) derived from dysregulation of flagellum growth between the two flagella during the cell cycle. This observation is consistent with Cep164C ablation causing continuous flagellum growth of the old flagellum over several cell cycles in the absence of a locking mechanism.

The ultrastructure of the flagellum and cell shape were analysed for abnormalities in case defective assembly would cause dysregulation. Scanning electron microscopy illustrated normal fusiform cell shape and form in cells, but with shorter and longer flagella compared to uninduced cells (Fig S1A-D). Transmission electron microscopy (TEM) did not reveal any visible abnormalities in the ultrastructure of the flagellum (Fig S1E,F) or basal bodies (Fig S1G). Longitudinal sections of the transition zone were analysed and measured. There were no observable abnormalities and there was no difference in the length of transition zones (Fig S1H-J; N=25 sections).

The grow-and-lock model predicts that a molecular lock is applied to prevent flagellum length change after assembly; locking the old flagellum into a stable configuration, but this is clearly absent in these induced RNAi cells. Importantly, since there are no abnormally long new flagella in dividing cells then the presence of abnormally long old flagella must occur via continued growth of the old flagellum in preceding cell cycles where the lock has not been applied (Fig 2K).

### Cep164C accumulates on locked flagella

During the cell cycle, the new flagellum grows up to 80% the length of the old one prior to cytokinesis, with final growth being completed before the next cell cycle [14]. During the next cell cycle, this flagellum becomes the old flagellum and does not increase in length or disassemble whilst an entirely new flagellum grows alongside. A locking mechanism is proposed to prevent the old flagellum from growing any longer or shortening in subsequent cell cycles [5]. Previous work showed that when the cell division inhibitor teniposide was applied for 8, 16 or 24hrs, the new flagellum grew to the same length as the old flagellum but never exceeded it. This illustrates an intrinsic locking event was initiated on the new flagellum to prevent further growth before cytokinesis [5]. We reasoned that if Cep164C was involved in the locking mechanism then Cep164C it could accumulate at the base of the new flagellum in these teniposide-treated cells once it had reached the length of the old. In order to test this, the published teniposide experiment was repeated, but this time using the endogenously tagged Cep164C:mNeonGreen cells at 8, 16 and 24hrs following addition of teniposide. In control cells, endogenously tagged Cep164C was only observed at the base of the old flagellum of dividing cells as expected (Fig 3A,B). In teniposide-treated cells, Cep164C was also found on the old flagellum in all dividing cells also as expected (Fig 3C; one signal), but, significantly, Cep164C also accumulated at the base of the new flagellum in ∼40% of cells (Fig 3D; two signals; N=200 cells), illustrating that Cep164C was indeed accumulating at the base of the new flagellum as well as the old flagellum in a proportion of cells.

Measurements of cells with old and new flagella were carried out in order to discover if accumulation of Cep164C onto the new flagellum in 40% of the teniposide-treated cells is correlated with the length of the new flagellum. In teniposide-treated cells with two Cep164C signals (old and new flagellum) the new new:old flagellum length ratio 84% and was almost identical to control post-mitotic cells at 82% (Fig 3E control late and ten+ 2 signals; N=200 cells). In teniposide-treated cells with only one Cep164C signal on the old flagellum the ratio of old:new flagellum length was only 42%, illustrating that ectopic Cep164C was not added to short new flagella. This was compared to G2 stage control cells where the new flagellum is still growing where there was a similar ratio of 46%. In summary, these experiments results support the published teniposide experiments [5] showing that there is a lock to new flagellum growth even in the absence of cytokinesis. These results show that endogenously tagged Cep164C accumulates at the base of new flagella once they are close to the length of the old flagellum. This is consistent with Cep164C being part of a locking mechanism.

### Appearance of ectopic Cep164C on the new flagella basal body

Given that loss of Cep164C leads to continuous flagellum growth on the old flagellum, we decided to test if ectopic expression of the protein leads to dysregulation of flagellum growth or if excess protein is targeted inappropriately to the new flagellum basal body. Inducible ectopic expression of Cep164C::mNG had no effect on growth of the population (SFig2A,B growth curve and western). Maximum expression was reached 24 hours after triggering induction. Inducible ectopically expressed Cep164C localised to the base of the old flagellum in all dividing cells with 2 flagella as expected (SFig 2D; OF). In addition, ectopically expressed Cep164C abnormally located to the base of the new flagellum as well as the old flagellum in 21% of dividing cells rising to 36% by 72hrs post-induction, albeit sometimes at a lower level than in the old flagellum (FigS2E; N=200 cells). Measurements of the new flagellum length in cells with Cep164C on the old flagellum only (one signal) or on the old and new flagellum (2 signals) revealed no significant difference in the mean average new flagellum length in these two populations. However, there was a significant variation in new flagellum lengths (SFig 2G; Supple. Table 1). This might be expected as Cep164C accumulation on the new flagellum appeared to be at a lower level, so the expression levels may not be effective in locking new flagellum growth. However, these results demonstrate that ectopically expressed Cep164C is specifically targeted to the old flagellum basal body and is also able to abnormally associate with the new flagellum basal body in some cells, which might occur when all the sites on the old flagellum for Cep164C basal body are occupied. Exactly how proteins are able to distinguish between flagella is still an unanswered question.

### Maintaining an old flagellum whilst growing a new flagellum

Our results demonstrate that a mature basal body only acquires Cep164C in the 3^rd^ cell cycle after initial formation and this coincides with completion of flagellum growth (Fig 4A). In subsequent cell cycles Cep164C is recruited to the old flagellum basal body and contributes to the locking mechanism by limiting further growth of the old flagellum (Fig 4B). However, maintenance of the old flagellum can still occur as intraflagellar transport (IFT) particles can still enter the old flagellum [5]. Knockdown of Cep164C by RNAi led to further growth of the old flagellum, but, significantly there was assembly of a shorter new flagella in the same cell. This indicates a dyregulation of growth between the old and new flagellum. One explanation could be that the new flagellum was unable to reach the normal length as material was also entering the old flagellum, suggesting an intrinsic limit of flagellum components within a cell. Certainly, it was been shown that the soluble pool of tubulin or other flagellar components (PFR) is low in Trypanosomes [15], but it is currently unknown whether there is a restricted access of tubulin to the locked old flagellum [5].

In mammalian cells, there is a single well characterised Cep164, which is critical for centriole docking via distal appendages and recruits the serine/threonine protein kinase TTBK2 to the centriole, which initiates a pathway for promoting the growth of cilia [9,11,16,17]. We have discovered an additional regulatory role in controlling ciliogenesis, possibly by limiting component access to an assembled flagellum. It appears that Cep164C could be part of a common regulatory mechanism in many ciliated unicellular organisms and is an under-appreciated mechanism of flagellum regulation.

There are many unicellular organisms that need to maintain existing flagella whilst growing new flagella during cell division, including *Paramecium, Giardia* and *Trichomonas sp.* to name a few [18]. Trypanosomes have three non-identical Cep164 proteins [19] with Cep164A and B localising to both old and new flagella [20]. Indeed, there are multiple Cep164 orthologues *in Paramecium, Tetrahymena, Giardia, Leishmania* and *Trichomonas* [19] indicating that Cep164 proteins potentially perform multiple functions in regulation of flagellum/cilium growth in eukaryotic organisms. We conclude that Cep164C is a molecular component of a lock to flagellum growth after assembly and is applied to the old flagellum basal body to prevent further growth.

## Acknowledgements

We would like to thank Prof. Keith Gull and Prof. Derrick Robinson for the kind gift of antibodies L8C4 and Mab25 used in this study. Dr Flavia Moreira-leite and Dr. Anna Frej for assistance. Dr Saad Arif for assistance with statistics. Work was funded by BBSRC BB/M000532/1 to SV. Research at the Institut Pasteur is funded by La Fondation pour la Recherche Médicale (Equipe FRM DEQ20150734356) and by a French Government Investissement d’Avenir programme, Laboratoire d’Excellence “Integrative Biology of Emerging Infectious Diseases” (ANR-10-LABX-62-IBEID). E.B. was supported by fellowships from French National Ministry for Research and Technology (doctoral school CDV515) and from La Fondation pour la Recherche Médicale (FDT20170436836).

---

**Supplemental Figure 1:**
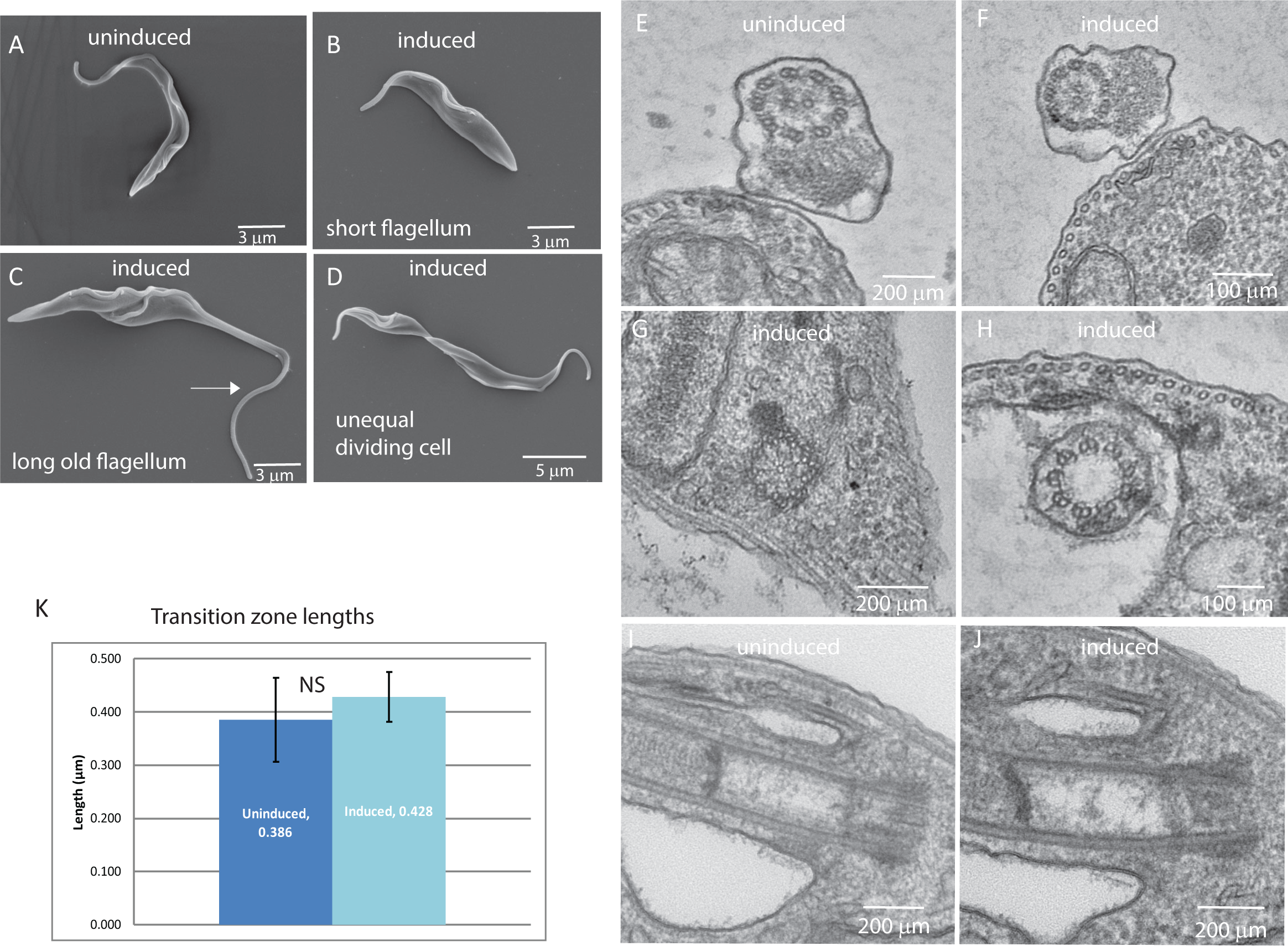
A-D: Scanning electron microscopy of uninduced (A) and induced *CEP164C*^*RNAi*^ cells (B-D) at 72hrs post-induction of RNAi illustrating small cells (B), dividing cells with a long old flagellum (C; arrow), dividing cell with one short cell and one long cell (D); E-J: transmission electron microscopy of *CEP164C*^*RNAi*^ cells. Cross section of the 9 + 2 microtubule axoneme in uninduced (E) and induced (F) cells, cross section of a basal body (G), cross section of induced transition zone, (H), longitudinal section of uninduced (I) and induced (J) transition zones; K: Measurements of transition zones in longitudinal TEM sections (N=25).

**Supplemental Figure 2:**
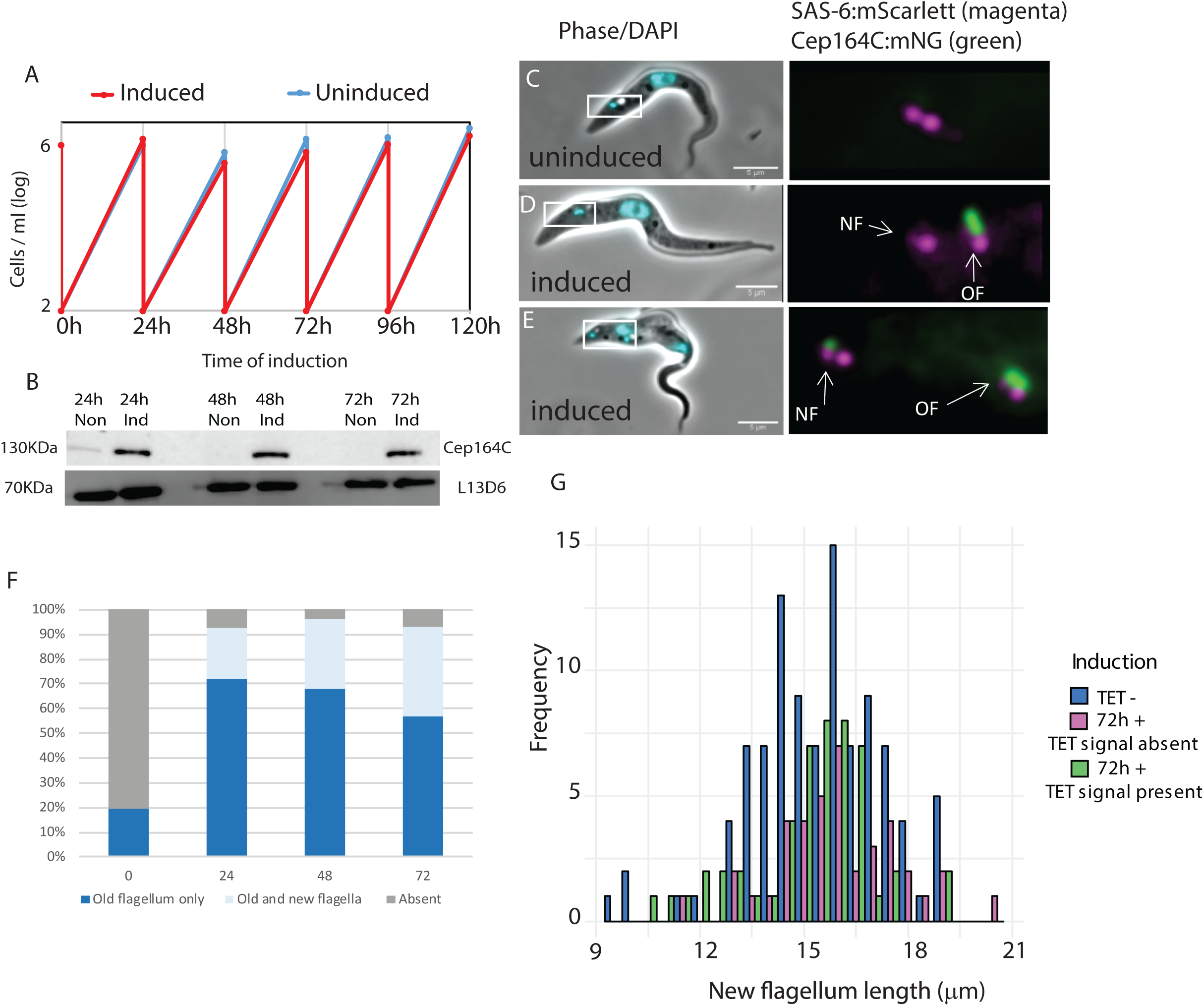
Inducible ectopic expression of Cep164C in endogenously tagged TbSAS-6 cells; A: Growth rate cells cultured between 2 × 10 ^6^ and 6 × 10^6^ cells per ml; B: Western blot using anti-Ty1 antibody BB2. This Ty1 epitope flanks the ectopic mNeonGreen Cep164C and is detected using Anti-Ty antibody BB2. Loading control using L13D6 antibody (flagellum marker); C-E localisation of ectopically expressed Cep164C::mNG (green) by live cell imaging in a cell line containing endogenously expressed TbSAS-6::mScarlet (magenta) as the basal body marker; C: Uninduced cells expressing endogenously tagged TbSAS-6:mScarlett only; D: induced dividing cell with Cep164C only on the old flagellum (OF; green); E: Induced cell with Cep164C on the new flagellum and old flagellum; F: Percentage of dividing cells with Cep164C on the old flagellum only (dark blue) or on the new and old flagellum of dividing cells (light blue) in 0h (uninduced), 24h, 48h, 72hs post-induction of ectopic Cep164C expression. Grey – no Cep164C signal; G: New flagellum length measurements in post-mitotic cells uninduced (blue – Tet -) and 72hrs post-induction (72h +). Ectopically expressed Cep164C present on old flagellum only and not on the new flagellum (72h + signal absent - magenta). Ectopically expressed Cep164C present on both the old and new flagellum (72h+ signal present – green) (N=200 cells).

**Supplemental Table 1:** Table of statistics for Cep164C experiments. 1F cells from Fig 2H, Cep164C dividing cells, Cep164C old flagellum (Fig 2I), new flagellum (Fig 2J); Fig 3F; Supplemental Fig 2G.

| **TbCep164c 1F Cells** | **Test-statistic (F)** | **P-value** | **df** | **Wilcoxon Test p-value** |
| --- | --- | --- | --- | --- |
| 24 vs. Uninduced | 0.21763 | 8.537e-25 | 200 | 9.2e-12 |
| 96 vs. Uninduced | 0.14378 | 2.3e-37 | 200 | 5.2e-08 |
| 96 vs. 24 | 0.66066 | 0.003618 | 200 | 0.5950663 |

**Fig 2 J Cep164C RNAi – 2F cells – New Flagellum**
| **Cep164C – New Flagellum** | **Test-statistic (F)** | **P-value** | **df** | **Wilcoxon Test p-value** |
| --- | --- | --- | --- | --- |
| 24 vs. Uninduced | 0.663677 | 0.004003 | 200 | 4.569e-51 |
| 96 vs. Uninduced | 0.6034171 | 0.000406 | 200 | 2.07439e-56 |
| 96 vs. 24 | 0.9092 | 0.503 | 200 | 0.03252 |

**Fig 2 I - Cep164C RNAi – 2F cells – Old Flagellum**
| **Cep164C – Old Flagellum** | **Test-statistic (F)** | **P-value** | **df** | **Wilcoxon Test p-value** |
| --- | --- | --- | --- | --- |
| 24 vs. Uninduced | 0.21307 | 2.170e-25 | 200 | 7.624e-11 |
| 96 vs. Uninduced | 0.12635 | 1.228e-41 | 200 | 2.024e-10 |
| 96 vs. 24 | 0.59299 | 0.0002515 | 200 | 0.4239183 |

**Fig 3 F – Old/new flagellum length ratios after teniposide treatment**
| **Old/new flagellum length ratios** | **Test-statistic (F)** | **P-value** | **Num df** | **Denom df** | **Wilcoxon Test p-value** |
| --- | --- | --- | --- | --- | --- |
| Control early vs control late | 4.457 | 7.362e-12 | 101 | 89 | 1.908e-31 |
| Control early vs teniposide 24h – one signal | 0.53393 | 0.0009785 | 101 | 139 | 0.166 |
| Control early vs teniposide 24h – two signals | 0.62546 | 0.02581 | 101 | 80 | 1.1396e-23 |
| Control late vs teniposide 24h – one signal | 0.1198 | 7.711e-22 | 89 | 139 | 2.363e-23 |
| Control late vs teniposide 24h – two signals | 0.14033 | 1.618e-17 | 89 | 80 | 0.0897 |
| Teniposide 24h – one signal vs teniposide 24h – two signal | 1.1714 | 0.4404182 | 139 | 80 | 4.958e-19 |

**New flagellum length**
| **New flagellum length** | **Test-statistic (F)** | **P-value** | **Num df** | **Denom df** | **Wilcoxon Test p-value** |
| --- | --- | --- | --- | --- | --- |
| 72h Signal Absent vs. TET- | 1.1808 | 0.567 | 100 | 40 | 0.1184 |
| 72h Signal Present vs TET- | 1.0827 | 0.7709 | 100 | 50 | 0.4548 |
| 72h Signal Absent vs<br>72h Signal Present | 0.91696 | 0.7856 | 40 | 50 | 0.04446 |

## Materials & Methods

### Parasites and culture conditions

SmOxP9 procyclic *T. brucei* were used for all experiments. These were derived from TREU 927 strain, expressing T7 RNA polymerase and tetracycline repressor [21]. Cells were cultured in SDM79 media supplemented with Hemin and 10% FCS [22] c at 28°C at a density between 1×10^6^ −1×10^7^ cells/mL.

### Endogenous tagging of Cep164C and immunofluorescence assay for co-localisation with RP2

For the endogenous tagging of Cep164C (Tb927.1.3560) with mNeonGreen (mNG), a PCR product was obtained using the oligos (5’-ATTTCTACGCTCACGCTATATTACTCTCTTTCTTTCCATTAGAAAGGTCGACTCATAGTCGAAATTTTTTTTTTGAAGCTGTATAATGCAGACCTGCTGCGTATAATGCAGACCTGCTGC −3’) and (5’-AGCCATTTCCCGTACTCCAGAAGCTCGGTTTCTGAAGGTTCGTAATTGTCGCTCGTCACAACATCTAGAACTACGGACATACTACCCGATCCTGATCCAG −3’), and the pPOTv6 (3Ty).mNG.blast as a template for the PCR reaction, and the transfection was carried out as previously described [23]. After clone selection, cytoskeleton preparations were made using these cells for use in immunofluorescence assays. Cells were washed twice with PBS (10 mM phosphate buffer, 2.7 mM, 137 mM sodium chloride, pH 7.2) by centrifugation at 800x g and settled onto glass slides for 5 minutes. The cell membrane was solubilized with 0.5% Nonidet P-40 in PEME (100 mM Pipes·NaOH (pH 6.9), 2 mM EGTA, 1 mM MgSO_4_ and 100 nM EDTA) for 30 s, followed by fixation in −20°C methanol for 20 min. Cytoskeletons were rehydrated in PBS and blocked with 1% BSA for 1 hour. After the blocking step, cells were incubated with anti-retinitis pigmentosa-2 (RP-2) (YL1/2 labels tyrosinated tubulin but also cross reacts with RP2) (1:40) [24] for 1 hour, followed by three 5 minute washes in PBS. A second incubation with anti-mouse IgG FITC (1:200) (Jackson ImmunoResearch) was carried out for 1 hour. Following secondary antibody incubation, the slides were washed three times in PBS, and Hoechst 33342 was used to stain DNA. The slides were mounted in 50 mM phosphate-buffered glycerol (pH 8.0). Mounted slides were observed and imaged by wide-field epifluorescence using a Zeiss Axioimager Z2 microscope fitted with a Hamamatsu ORCA-Flash 4.0 camera and a 100× oil immersion objective with a numerical aperture (NA) of 1.46.

### Cep164C RNAi cell line construction and PFR labelling

RNAi plasmid for the knockdown of Cep164C was generated using the pQuadra system [25]. For this, oligos were designed for the amplification of Cep164C using RNAit [26] (Forward: 5’-ataccaatgtgatggTCTGAAACGCATTTGTCTGC-3’ and reverse: 5’-ataccatagagttggGCACAGCACAGAGGTTGAAA-3’). These oligos were used in PCR reactions using *T. brucei* genomic DNA as a template. The PCR product and the pQuadra 1 and pQuadra 3 plasmids were ligated, cloned and sequenced. Following the sequencing confirmation, the plasmid was digested with NotI (New England Biolabs) at 37° C for 16 hours and transfected into the endogenously tagged Cep164C:mNG cell line as previously described. After clone selection, stable cell lines were induced by addition of doxycycline (1µg/mL) to express dsRNA and consequent knockdown of Cep164C expression. Cell growth rate was monitored using a Beckman Coulter counter. Cells were collected at 0, 24, 48, 72 and 96 hours post-induction for immunofluorescence and western blotting assays. The immunofluorescence assays were performed as described in the previous topic. The monoclonal antibody L8C4 (1:200) (kind gift from Prof. Keith Gull), which recognises the paraflagellar rod [27] was used as a flagellum marker.

### Western blotting

Induced Cep164C cells were boiled at 100°C in Laemmli loading buffer (4% SDS, 20% glycerol, 0.12M Tris-HCl pH 6.8, 0.2 bromophenol blue) 5 minutes. Protein from 1×10^6^ cells was separated on a 10% SDS-PAGE gel, followed by a transfer to a PVDF membrane using the Mini Trans-Blot system (BIORAD), according to the manufacturer’s protocol. The transferred membrane was blocked with 5% skimmed milk (Sigma) in 0.1% TBS-Tween for one hour, and then incubated with the anti-TY (1:200) monoclonal mouse antibody for one hour. Three membrane washes were performed with TBS-tween (Tris 50mM, NaCl 100 mM, pH 7.6, tween 0.1%) and incubated with anti-mouse IgG coupled to horseradish perixodase (Jackson), diluted 1:10,000, for 1 hour. The membrane was washed three times with TBS-tween following secondary antibody incubation. For final detection of protein, the chemiluminescence reaction was performed with WesternBright Quantum HRP substrate (Advansta) according to the manufacturer’s protocol. L8C4 was used as a loading control. The western blot assay to confirm the ectopic expression of Cep164C was performed as described above with the addition of the L13D6 antibody (1:100) as a loading control.

### Teniposide treatment

Teniposide (Sigma SML0609), a topoisomerase II inhibitor of cell division, was dissolved in DMSO and added to 1.5×10^6^ cell/ml culture of the endogenously tagged Cep164C:mNeonGreen cell line at a final concentration of 200 μM. In the control flask, the same volume of DMSO without teniposide was added. Samples were collected at 0, 8, 16 and 24 hours after teniposide treatment. Immunofluorescence of extracted cytoskeletons was carried out as described above. The monoclonal antibody mAb25 (1:20) (kind gift from Prof. Derrick Robinson), which recognises the axonemal protein TbSAXO1 [28] was used as flagellum marker, and anti-mouse IgG TRITC (1:200) (Jackson ImmunoResearch) was used as secondary antibody.

### Transmission and scanning electron microscopy

Non-induced control and 96h-induced Cep164C RNAi cell line samples were collected and prepared for thin-section transmission electron microscopy analysis as previously described [29]. 70nm thin sections were made by PTPC: PowerTome PC (RMC Boeckeler) and stained using lead citrate for five minutes, followed by three washes with MiliQ water. Images were captured using a Hitachi H-7650 transmission electron microscope

For scanning electron microscopy, non-induced control and 96-h induced Cep164C samples were collected and prepared as previously described [29]. Images were captured using a Hitachi S-3400 microscope.

### Inducible Cep164C ectopic expression assay and live cells imaging

For ectopic expression of Cep164C::mNG, oligos were designed (Forward 5’-ttaactagtATGTCCGTAGTTCTAGATG-3’ and reverse 5’-aaaggatccTTAATGAGCCGATTTACCC-3’) for the amplification of the gene, using *T. brucei* genomic DNA as a template. The PCR product was cloned into the pDEX-777 [21] plasmid, which was transfected into a *T. brucei* SmOx p927 cell line with TbSAS-6:mScarlet endogenously tagged at the N-terminus. After clone selection with 5µg/mL phleomycin, the cells were prepared for live cell imaging. Cells were first washed by centrifugation at 800× *g*, followed by re-suspension in PBS supplemented with 10 mM glucose and 46 mM sucrose (vPBS). In the second wash, DNA was stained using 10 µg/mL Hoechst 33342. After the third wash, cells were re-suspended in vPBS and settled onto glass slides. The ectopic expression of Cep164C:mNG and endogenous expression of SAS6:mScarlet was visualised using an epifluorescence microscope as described previously.

### Flagellum measurements

Measurements of flagellum length were carried out using ImageJ on images of cytoskeletons labelled with either anti-axoneme antibody mAb-25 or an anti-PFR antibody L8C4.

### Statistics and graphs

The statistical tests, F-Test and Wilcoxon, were performed in RStudio. Graphs for the analysis were made in RStudio and Excel 2016 (Microsoft).

